# Mitochondrial instability contributes to IFN-driven heart disease

**DOI:** 10.1101/2025.09.15.674036

**Authors:** Arati Naveen Kumar, Amelia Meecham, Deepika Rai, Brennan Kandalaft, Trinitee Oliver, Aleksandr B. Stotland, Asli E. Atici, Magali Novas Rivas, Richard Ainsworth, Megan L. McCain, Sarah J. Parker, Caroline A. Jefferies

## Abstract

Type I interferons (IFNs) are linked to an increased risk of cardiovascular disease and are chronically elevated in systemic autoimmune diseases such as systemic lupus erythematosus (SLE). We evaluated the effect of chronic IFNα exposure on heart function using the resiquimod model of IFN-inducible SLE. Echocardiography showed that C57BL/6 mice had reduced left ventricular function following administration of resiquimod (R848) for 4 weeks. This was accompanied by increased α-SMA deposition, infiltration of both innate and adaptive immune cells and a robust increase in fibroblasts. Importantly endothelial cell abundance decreased, as measured by CD31 staining. Transcriptomic and proteomic analysis showed a concordant increase in pathways representing type I IFN response, ischemia, oxidative stress, fibrosis and cardiomyocyte stress, whereas genes and proteins representing oxidative phosphorylation were decreased, as were metabolites on the TCA cycle, suggesting impaired ATP production in the R848-treated hearts. As cardiomyocytes are the most energetically demanding cell type in the heart, and the cells most susceptible to stress and inflammation, we assessed the effects of chronic IFN exposure on their function. Chronically exposed AC16 cells showed impaired basal respiration, increased mitochondrial depolarization, increased mitochondrial mass and fragmentation, suggesting mitochondrial stress. mtDNA release is a key consequence of mitochondrial stress, driving cGAS activation and induction of IFN-stimulated genes. Not only was enhanced mtDNA detectable in chronically activated AC16s, but the increased expression of ISGs could be inhibited by the cGAS inhibitor G140. Thus chronic IFN exposure promotes cardiomyocyte stress, mitochondrial DNA release and a feed forward inflammatory loop driven by cGAS, a recently identified mediator of cardiovascular disease. Importantly, human iPSC-derived cardiomyocytes recapitulated our key findings in AC16 cells, and cGAS inhibition could rescue ISG expression and mitochondrial depolarization.

## Introduction

Heart disease is a significant and well-recognized complication of systemic lupus erythematosus (SLE), contributing substantially to increased morbidity and mortality in affected individuals (1). Patients with SLE have a markedly elevated risk of premature and accelerated atherosclerosis, resulting in heightened incidence of ischemic heart disease and myocardial infarction even at a young age (2). In addition to atherosclerosis, lupus can also cause myocarditis, pericarditis, valvular heart disease, and coronary microvascular dysfunction (CMD), reflecting the multifaceted ways autoimmunity and chronic inflammation can injure the heart (3). Many immune features of SLE such as elevated type I interferons, chronic inflammation, and persistent autoantibody production are strongly correlated with increased cardiovascular risk, but how they contribute to increased cardiovascular disease (CVD) risk in SLE is currently unknown.

Type I interferons (IFN) play key roles not only in host defense against infections but also in the development and progression of various forms of heart disease. IFN signaling can be triggered by cardiac injury, ischemia, and viral infection, leading to activation of hundreds of interferon-stimulated genes (ISGs) and substantial shifts in cardiac immune and metabolic environments (4). In the heart, induction of the type I IFN response has been reported in ischemic and non-ischemic cardiomyopathy, as well as in viral myocarditis (5). The cardiac effects of interferons are context-dependent: while they are protective in viral myocarditis, in the absence of infection (such as ischemia or in pressure overload), persistent IFN signaling can promote adverse cardiac remodeling, ultimately accelerating heart failure (6-8). Our previous analysis of genes differentially expressed in whole blood RNA samples from SLE patients with coronary microvascular dysfunction (CMD), highlighted differences in IFN signaling, cytosolic RNA/DNA sensing pathways between patients with CMD versus those without (9).

As the primary source of ATP in cardiomyocytes, mitochondria are essential for maintaining contractile function and cellular viability; their impairment disrupts energy production, leading to reduced cardiac output and myocyte death (10). In heart disease, mitochondrial dysfunction has emerged as a key pathogenic factor, underlying conditions such as heart failure, ischemic cardiomyopathy, and diabetic cardiomyopathy (10). Damaged mitochondria exhibit impaired oxidative phosphorylation, diminished ATP generation, and increased production of mitochondrial reactive oxygen species (mtROS), which drive oxidative stress and proteotoxic injury due to misfolded proteins in cardiomyocytes (11). Excessive mtROS activates signaling pathways that induce cell death, inflammation, and adverse cardiac remodeling, exacerbating cardiac dysfunction. Compromised mitochondrial dynamics such as imbalances in fusion, fission, and defective mitophagy lead to the accumulation of dysfunctional mitochondrial fragments, disrupt energy supply, and further contribute to myocardial injury (12). These maladaptive shifts in mitochondrial dynamics disrupt myocardial energetics and cellular survival mechanisms, promoting cardiac remodeling and the progression of heart failure (10). An additional consequence of mitochondrial stress is the release of mitochondrial DNA (mtDNA) into the cytoplasm, engaging cytosolic DNA sensors, such as cyclic GMP-AMP synthase (cGAS), and downstream activation of the stimulator of interferon genes (STING) pathway (13). mtDNA and mitochondrial DAMPs additionally activate toll-like receptors (TLRs) in plasmacytoid dendritic cells, setting off cascades of pro-inflammatory cytokine release, including IFN-α (13). Our previous work has shown that chronic IFN exposure of monocytes induces mitochondrial reprogramming and epigenetic reprogramming of the IFN response (14). However, whether chronic IFN exposure triggers similar changes in the heart remains unknown.

In this study we investigate the effects of chronic activation of TLR7/8 in a mouse model of IFN-driven SLE-like disease on cardiac function. Using this model we show that chronic IFN induction in mice results in defective heart function, increased representation of pathways contributing to cardiac stress (ischemia/hypoxia, cardiomyocyte remodeling, oxidative stress) and a decrease in those regulating mitochondrial function and ATP production. Addressing how chronic IFN affects cardiomyocytes, the most ATP-dependent cell type in the heart, we find that chronic IFN induces proteomic and transcriptomic changes in AC16 cardiomyocytes consistent with inflammation and altered mitochondrial function. Mitochondrial changes are accompanied by increased mitochondrial fragmentation and release of mtDNA into the cytoplasm and ISG expression, thus contributing to a feed forward augmentation of chronic IFN effects. Importantly, targeting cGAS can reverse these effects. Thus, targeting cytosolic RNA/DNA sensing and the type I IFN response may reduce CVD risk in SLE patients.

## Materials and Methods

### Animals

Ethical Compliance: All procedures followed National Institutes of Health guidelines and received approval from the Cedars-Sinai Medical Center Institutional Animal Care and Use Committee under protocol IACUC-008858. Both male and female C57BL/6J wild-type mice (9-12 weeks old) underwent epicutaneous administration of 100 μg of Resiquimod dissolved in 30 μl of a 1:2 ethanol/acetone mixture. The solution was applied topically to the left ear three times weekly over a four-week period. Echocardiograms were performed on all animals, which were subsequently sacrificed at the study endpoint for organ and blood collection.

### Cell line

AC16 human ventricular cardiomyocytes were maintained and utilized for experiments at passages 4 through 10 in 6-well plates (100,000 cells per well). The cells were cultured for two days in growth medium (DMEM/F12 supplemented with 12.5% FBS, 1% antibiotic-antimycotic, pH 7.4) until they reached 70–75% confluence.

### In-vitro model

AC16 cardiomyocytes were seeded at 100,000 cells per well in complete DMEM and initially stimulated for 24 hours with IFNα (1000 U/ml) or PBS. After stimulation, the cells were washed and rested for 4–5 days. They were then washed again, replated at 100,000 cells per well, and subjected to a second 24-hour stimulation with IFNα (1000 U/ml) or PBS. Following this, cells were washed and collected for RNA and protein extraction.

### Real-time Quantitative Polymerase Chain Reaction (RT-qPCR)

RNA was isolated from tissues and cultured cells with TRIzol reagent (Sigma-Aldrich) following the manufacturer’s instructions. Reverse transcription was performed, and quantitative real-time PCR results were analyzed using the ΔΔCt method, normalized to 18s rRNA expression.

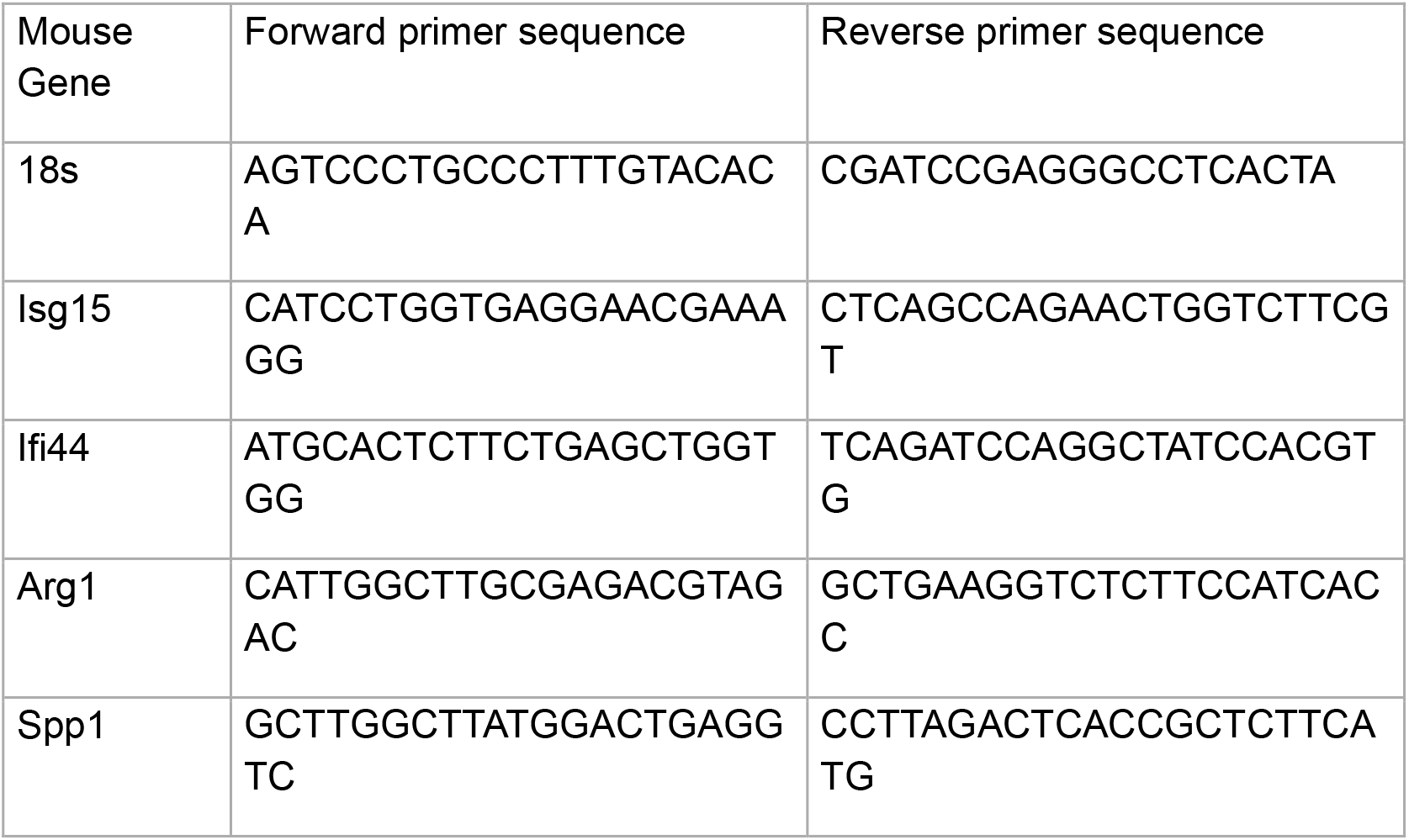
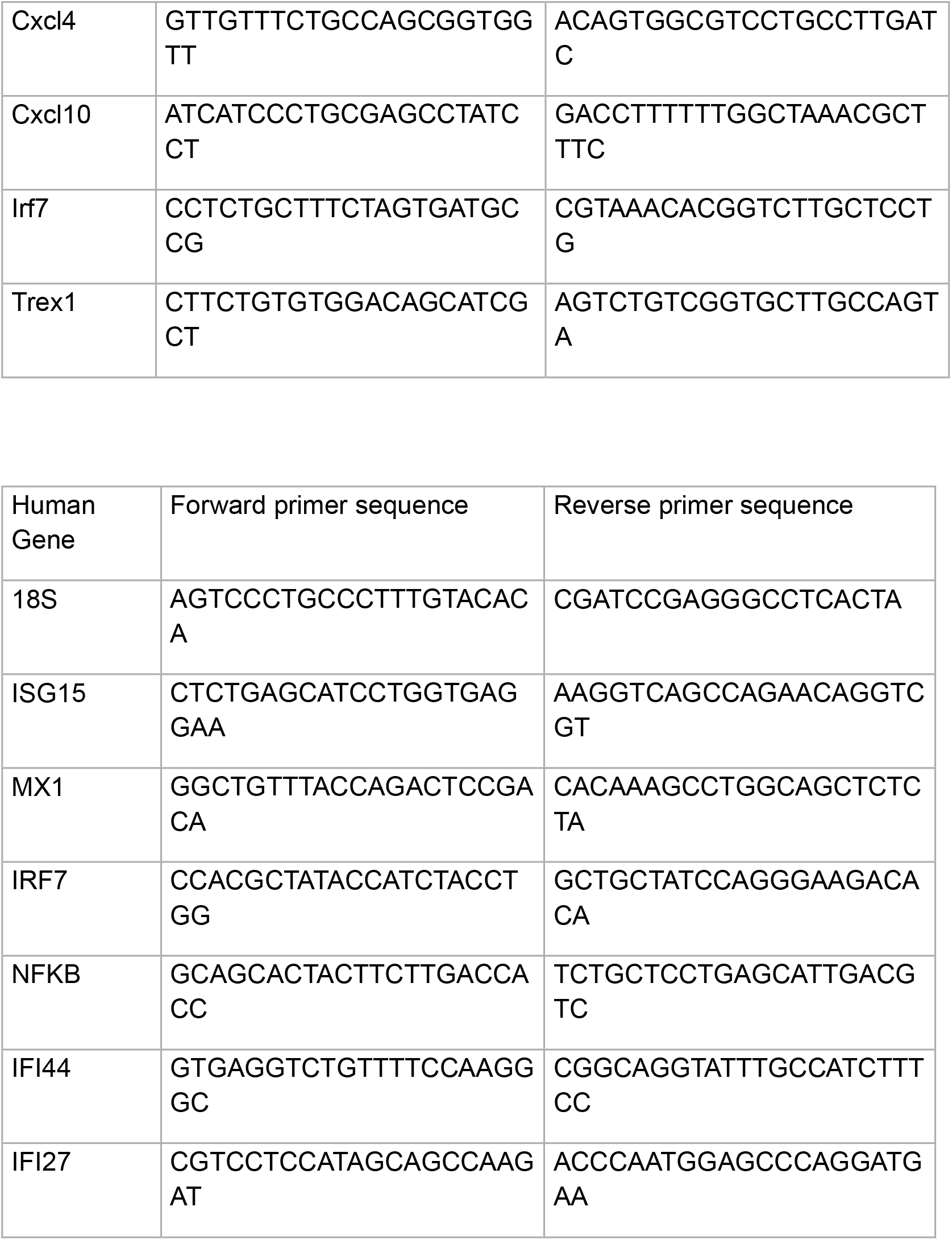

### mRNA-Seq library preparation and sequencing

RNA integrity was assessed on a 2100 Bioanalyzer using the Agilent RNA 6000 Nano Kit (Agilent Technologies), and RNA quantification was performed with the Qubit fluorometer and Qubit RNA HS Assay Kit (ThermoFisher Scientific). A total of 325 ng RNA per sample underwent mRNA purification using the NEBNext® Poly(A) mRNA Magnetic Isolation Module (New England Biolab Inc). Stranded RNA-Seq libraries were constructed with the IDT xGen Broad-Range RNA Library Prep Kit (Integrated DNA Technologies) and amplified by 12 PCR cycles. Library concentrations were determined with the Qubit 1X dsDNA HS Assay kit, and library sizes evaluated using the Agilent 4200 TapeStation with Agilent HS D1000 ScreenTape. Multiplexed libraries were pooled for sequencing on a NovaSeq X (Illumina) platform using single-end 75 bp reads at a depth of 30 million reads per sample.

### Bioinformatics and data analysis

Raw reads obtained from RNA-Seq were aligned to the transcriptome using STAR (version 2.5.0)/RSEM (version 1.2.25) with default parameters, using a custom human GRCh38 transcriptome reference downloaded from https://www.gencodegenes.org, containing all protein coding and long non-coding RNA genes based on human GENCODE version 33 annotation. Expression counts for each gene in all samples were normalized by a modified trimmed mean of the M-values normalization method and the unsupervised principal component analysis (PCA) was performed with DESeq2 Bioconductor package version 1.42.0 in R version 4.3. Each gene was fitted into a negative binomial generalized linear model, and the Wald test was applied to assess the differential expressions between two sample groups by DESeq2. Benjamini and Hochberg procedure was applied to adjust for multiple hypothesis testing, and differential expression gene candidates were selected with a false discovery rate less than 0.05. For functional enrichment analysis across sample groups, we conducted genes enrichment analysis using the R package “clusterProfiler v4.10.0” (15) and “pathfindR v2.3.1 (16). Data is included in supplementary data files 1-3. Gene set enrichment analysis (GSEA) was performed against the Reactome pathway database (version v2025.1) from MSigDB. Significance was assessed using 1,000 permutations, and the top-ranking pathways were identified based on the normalized enrichment score (NES) and a false discovery rate (FDR) q-value < 0.25. Venn diagrams were compiled from differentially expressed genes and proteins using Venny 2.1.0.

### Protein extraction from pellets and Western Blot analysis

Protein extraction from cell pellets was done by resuspending pellets in ice-cold RIPA buffer supplemented with a protease and phosphatase inhibitor cocktail to prevent protein degradation and preserve post-translational modifications. Samples were incubated on ice for 30 minutes with intermittent vortexing to ensure thorough lysis, followed by centrifugation at 14,000 x g for 15 minutes at 4°C. The clarified supernatant containing solubilized proteins was collected for downstream quantification and analyses. For western blot analysis proteins were separated on 7% to 12% Tris-acetate gel (BioRad) and electrophoretically transferred onto PVDF membranes and incubated with primary anti-bodies (

### Mitoplex

Tier 2 targeted proteomic analysis, as previously described by Stotland et al, was employed to quantify mitochondrial proteins. Experimenters were aware of group assignments during sample processing and analysis. Adipose tissue was lysed in 8 M urea with 1 M ammonium bicarbonate (pH 8.0), and protein concentration was determined using the Pierce BCA assay (Thermo Fisher Scientific). Samples were digested using the SP3 protocol: 20 μg of protein was adjusted to 60 μl with 6 M urea, 1 M ammonium bicarbonate, and 5% SDS lysis buffer. Reduction was performed with 16.8 μl of 200 mM dithiothreitol for 30 min at 37°C and 300 rpm, followed by alkylation with

21.2 μl of 400 mM IAA for 30 min in the dark at room temperature. Tris-HCl (pH 8) was added to reach 160 μl total volume, then 5 μl of a 10:1 mass ratio bead suspension (1:1 hydrophilic/hydrophobic beads, Cytiva) was added and vortexed. Samples were adjusted to 70% ACN and incubated for 18 min, followed by on-magnet solvent removal and rinses with 2× 80% ethanol and 2× ACN (200 μl each). The dried sample was suspended in 50 mM Tris-HCl (pH 8) with 10 mM CaCl2 and digested with trypsin at a 1:20 ratio. Samples underwent 5 min bath sonication before overnight incubation (18 h, 37°C, 1200 rpm). After digestion, beads were removed, and samples were brought to 0.1% FA and 2% DMSO, then spiked with a 1:250 dilution of stable isotope-labeled reference peptides as described in Stotland et al. Eight μg of digested peptides, run in duplicate technical replicates, were separated on a Shimadzu Prominence UFLCXR HPLC using a Waters Xbridge BEH30 C18 column (2.1 mm × 100 mm, 3.5 μm, 0.25 ml/min, 36°C) coupled to a SCIEX QTRAP 6500. Mobile phase A contained 2% acetonitrile, 98% water, and 0.1% formic acid; B was 95% acetonitrile, 5% water, and 0.1% formic acid. After loading and equilibrating at 5% B and eluting peptides with a linear 5–35% gradient over 30 min, the column was washed with 98% B for 10 min, returning to 5% for re-equilibration. A scheduled acquisition method was used, monitoring peptide fragments within a 2-min retention window. Raw data were processed in Skyline (version 21.1.0.146) to select peak boundaries and quantify peak areas. Peaks were manually inspected, and unusable data excluded. Automated peak integration was applied, and processed data were exported for further analysis in R (Stotland et al). Only fragments with a CV ≤ 20% across heavy standards were quantified. Endogenous-to-heavy standard abundance ratios were averaged to generate final protein quantification values. Technical replicate abundance ratios were calculated per sample. Statistical analysis was performed in MetaboAnalyst 6.0 (http://www.metaboanalyst.ca).

### Metabolomic Analyses (dMRM)

Metabolites were extracted from frozen tissues using 800 μl of extraction buffer (80% methanol, 20% water, 0.8% ammonium bicarbonate) and homogenized on ice with a 5-second pulse at 50% power (Polytron PowerGen 125, Thermo Fisher Scientific). Samples were centrifuged at 21,000g for 10 min at 4°C, and supernatants were vacuum dried. Metabolite pellets were resuspended in methanol:water (20:80) for analysis on an Agilent 6470A triple quadrupole mass spectrometer (operated in negative mode) with an Agilent 1290 UHPLC and MassHunter dMRM database/method, detecting 219 polar metabolites per sample. The method utilizes ion-pair reversed-phase chromatography (IP-RP) for anionic/hydrophobic separation, with TBA as the ion pair reagent. Mobile phase A is water with 3% methanol, 10 mM TBA, 15 mM acetic acid. Phases B and D are isopropanol and acetonitrile, respectively; C is methanol with TBA and acetic acid. Separation was performed on an Agilent ZORBAX RRHD Extend-C18 (2.1 × 150 mm, 1.8 μm) with a matching guard column. The method leverages known retention times for dynamic MRM transitions. QQQ chromatograms were inspected in Agilent MassHunter Quantitative Analysis, and peaks manually checked. Data were analyzed in MetaboAnalyst 6.0.

### LC-MS/MS Analysis

DIA analysis for heart and AC16 cell proteins was performed on an Orbitrap Ascend Tribrid (Thermo Scientific) mass spectrometer interfaced with a EASY-Spray™ ionization source (Thermo Scientific, ES081) coupled to Vanquish Neo ultra-high-pressure chromatography system with 0.1% formic acid in water as mobile phase A and 0.1% formic acid in acetonitrile as mobile phase B. Peptides were separated at constant flow rate of 1.20 µL/minute with a linearly increasing gradient of 8-34% B for 0-90 minutes, then flushed with 98% B from 90-112 minutes. The column used was Thermo Scientific™ µPac™ HPLC column with a 200cm bed length (P/N: COL-NANO200G1B). MS1 resolution was set to 120,000 with AGC target set to standard. RF Lens was set to 60% with a maximum injection time of 251 ms. Fragmented ions were detected across a scan range of 380-985 m/z with 60 non-overlapping data independent acquisition precursor windows of size 10 Da. MS2 resolution was set to 15,000 with a scan range of 145-1450 m/z, a normalized collision energy of 30%, and a normalized AGC target of 400% with a custom maximum injection time set to 25 ms. All data is acquired in profile mode using positive polarity.

DIA analysis for heart and AC16 proteins was performed on an Orbitrap Astral (Thermo Scientific) mass spectrometer interfaced with an EASY-Spray™ nano-electrospray ionization source (Thermo Scientific, ES081) coupled to Vanquish Neo ultra-high-pressure chromatography system with 0.1% formic acid in water as mobile phase A and 0.1% formic acid in acetonitrile as mobile phase B. Peptides were separated at an initial flow rate of 3 µL/minute and 4% B for 0.5 minutes. At 0.5 minutes, the flow rate is decreased to 1.3 µL/minute and the gradient was sharply increased linearly from 4-9% B for 0.5-0.6 minutes. At 0.6 minutes, the flow rate was again decreased to 0.8 µL/minute and the gradient was increased from 8-22.5% B for 0.6-13.9 minutes. At 13.9 minutes, the gradient was ramped up slightly faster, going from 22.5-35% B from 13.9-20.8 minutes. At 20.8 minutes, the flow rate was increased to 2 µL/minute and the gradient was linearly increased from 35-55% B from 20.8 minutes to 21.2 minutes. At 21.2 minutes, the column undergoes a wash of 99% B with a flow rate of 3 µL/minute before being equilibrated for the next run. The column used was PepSep C18 15cm × 150 µm, 1.9µm (Bruker, P/N: 1893471). Source parameters were set to a voltage of 2000 V and a capillary temperature of 280°C. MS1 scan range was set to 380-980 m/z and MS1 resolution was set to 240,000 with an AGC target set to ‘Custom’ and a normalized AGC target set to 500%. RF Lens was set to 40% with maximum injection time set to 5ms. Precursor mass range was set to 380-980 m/z with 199 non-overlapping data independent acquisition precursor windows of size 3 m/z. MS2 scan range was set to 100-1000 m/z and normalized HCD collision energy set to 25%. Maximum injection time was set to 5ms with AGC target set to ‘Custom’ and normalized AGC target set to 500%. All data is acquired in profile mode using positive polarity.

### Data Integration

MetaboAnalyst 6.0 was used to integrate transcriptomic and metabolomic data in our study. Network-based multiomics data integration was applied to differentially expressed genes and metabolites using GO bP pathway library for annotation. Both metabolic and regulatory pathways identified were significantly altered (FDR q<0.02). The pathway impact value was also determined. An impact score 0.3 was used as cut-off.

### α-SMA immunostaining and quantification

Immunofluorescence staining of cardiac tissue sections was performed as previously described [reference]. Briefly, tissue sections underwent citrate-based heat-induced antigen retrieval, followed by immunostaining for α-smooth muscle actin (α-SMA; [manufacturer, catalogue number]). Stained tissue sections were analyzed using QuPath software. Myocardial regions of interest were manually annotated, excluding large vessels and non-tissue regions. α-SMA-positive staining was quantified using Gaussian smoothing (σ = 1) and a consistent fluorescence intensity threshold of 400 at a pixel resolution of 0.69 µm/pixel. Results were expressed as the percentage of α-SMA-positive area relative to the total analyzed myocardial area.

### Mitochondrial membrane potential analysis

Mitochondrial membrane potential (ΔΨm) was assessed in AC16 cardiomyocytes using JC-1 dye (Invitrogen; [catalogue number]). Following the indicated treatments, cells were incubated with JC-1 (2 µM) for 20 minutes at 37°C, protected from light. Fluorescence was measured using a Sony ID7000 spectral flow cytometer. Mitochondrial membrane potential was assessed by calculating the ratio of red JC-1 aggregate fluorescence to green JC-1 monomer fluorescence.

### Mitochondrial and lysosomal imaging and analysis

Following the indicated treatments, AC16 cardiomyocytes were stained with MitoTracker Green FM (MedChemExpress, HY-135056) and LysoTracker Red DND-99 (Invitrogen, L7528), followed by nuclear counterstaining with DAPI. Fluorescence images were acquired using an Echo Revolve microscope equipped with a 40× oil-immersion objective. Mitochondrial morphology was assessed through analysis of mitochondrial network organization and branching, while mitochondrial fluorescence intensity was quantified relative to nuclear staining. Lysosomal staining was assessed using fluorescence-based image analysis. Consistent image acquisition and analysis parameters were applied across experimental groups.

### mtDNA detection

Mitochondrial DNA (mtDNA) was isolated from cytosolic fraction of the cell samples using the Qiagen DNeasy Blood & Tissue Kit according to the manufacturer’s protocol. mtDNA collected was used for PCR to measure mitochondrial and housekeeping DNA coding regions.

### Oxygen Consumption and Acidification Analysis

AC16 human ventricular cardiomyocytes were analyzed via real-time respirometry using a Seahorse XFe96 Analyzer (Seahorse Biosciences), after IFNα stimulation. Oxygen consumption rate (OCR) served as a measure of oxidative phosphorylation and Extracellular Acidification Rate (ECAR) primarily measures the rate of the glycolytic pathway. They were measured using the Mito Stress and Glycolytic Stress Tests respectively. Purified AC16 cells were washed with 10 ml DPBS, resuspended in XF RPMI medium (Agilent Technologies), and plated at 10,000 cells/well in 200 μl on XF96 plates and equilibrated at 37°C for 1 hour before analysis.

Final concentrations of inhibitors for MitoStress assay were: 0.5 µM FCCP and 0.5 µM/0.5 µM antimycin/rotenone.

Final concentrations of inhibitors for Glycolytic Stress Assay were: Glucose: 10 mM, Oligomycin: 1 μM, 2-deoxy-glucose (2-DG): 50 mM

### Statistical Analysis

Data are shown as mean ± SD. Student’s unpaired t-test or one-way ANOVA with Bonferroni post-hoc testing were used for significance, with normality checked by Shapiro–Wilk test. Non-normally distributed data were analyzed by Mann–Whitney test or Kruskal–Wallis ANOVA with Dunn’s post-hoc test. All analyses were conducted in Prism v7.0a (GraphPad). Significance was defined as p < 0.05. Figure legend asterisks: *p < 0.05; **p < 0.01; ***p < 0.001; ****p < 0.0001.

## Results

Cardiovascular complications and coronary microvascular disease are a signficant comorbidity in SLE. To study the role of chronic IFN exposure in driving SLE CVD, we adopted the resiquimod (R848) model of IFN-inducible SLE, which had been previously shown to drive severe dilated cardiomyopathy and histopathology revealed widespread heart inflammation resembling autoimmune pancarditis in the recombinant inbred CFN hybrid mice (cross of C57Bl6/J, FVB/NJ, and NOD/ShiLtJ) (17). In our study we used C57Bl/6 mice, treating them with 100µg of R848 or vehicle control, 3 times a week for 4 weeks. At the end of the 4 weeks, echocardiography was performed to determine heart function, after which the mice were euthanized and tissues harvested for analysis. As expected, R848-treated C57BL/6 mice had significantly larger spleens than vehicle-treated controls and increased expression of IFN stimulated genes in splenocytes (Supplemental Figure 1A - C). Hematoxylin & eosin staining of paraffin-embedded heart sections showed a significant increase in left ventricular wall thickness and increased nuclei per um^2^ of tissue when compared to that of the control mice, indicating cellular hypertrophy (Figure 1A). Echocardiography showed that fractional shortening, ejection fraction, left ventricular posterior wall in systole (LVPWs) and cardiac output were all decreased (Figure 1B), consistent with left ventricular systolic dysfunction. Consistent with a loss of LV systolic dysfunction, we observed increased levels of α-SMA in R848 treated hearts, in addition to increased representation of transcripts associated fibroblasts (using the murine Microenvironment Cell Population counter R package), both indicative of increased fibrosis (Figure 1C). This was also accompanied by infiltration of immune cells including neutrophils, monocytes, macrophages and CD3 positive T cells (Supplemental Figure 2). As prolonged exposure to type I IFNs can negatively impact heart function by driving vascular remodeling and localized hypoxia, we assessed capillary density in the heart muscle of resiquimod treated mice. As shown in Figure 1D, resiquimod treatment significantly reduced the number of CD31+ cells/nucleus in the heart, as well as transcripts associated with endothelial cells (using the murine Microenvironment Cell Population counter R package), demonstrating a reduction in capillary density as a result of resiquimod treatment, corresponding to ischemic changes following R848 treatment. Thus, R848 treatment of C57BL/6 mice results in cardiac dysfunction, increased fibrosis and endothelial cell pruning.

**Figure 1.**
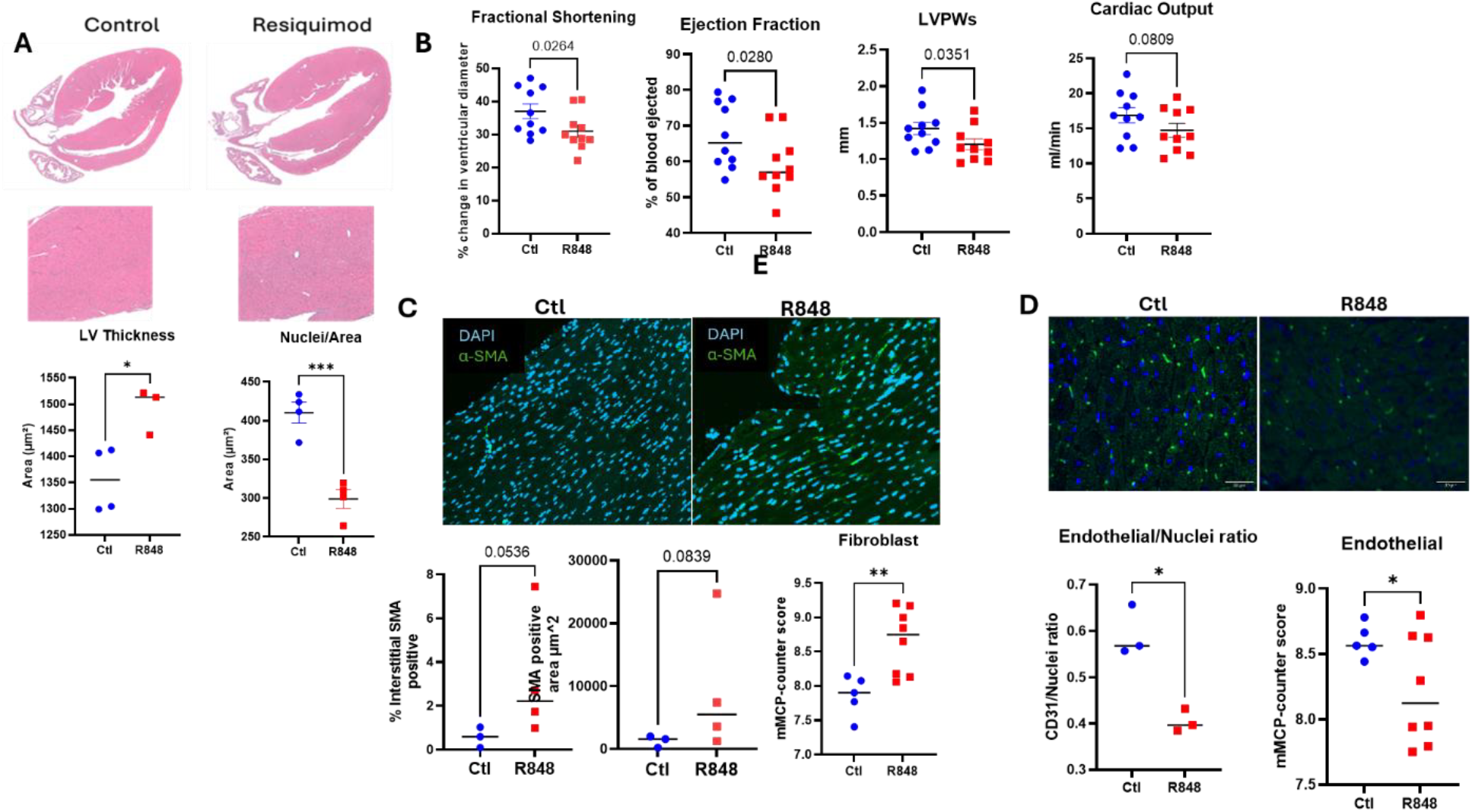
Resiquimod (R848) treatment induces cardiac inflammation, fibrosis and reduces cardiac function. (A) Paraffin-embedded heart sections from control (n=9) and R848-treated (n=10) mice were stained with Hematoxylin & eosin staining and the left ventricular (LV) thickness, number of nuclei per µM^2^ of tissue and LV lumen area were measured; (B)Echocardiograms were performed on control and R848-treated mice at 4-weeks 1 day prior to sacrifice and fractional shortening, ejection fraction percentage, left ventricular posterior wall shortening (LVPWs) and cardiac output measured; (C) Heart sections were stained for anti-a-SMA, images acquired at 10x magnification and quantified using ImageJ analysis; changes in fibroblast levels in the heart were determined from bulk RNAseq analysis using murine Microenvironment Cell Population counter (mMCP-counter); (D) The degree of vascularization was measured by comparing the degree of CD31 staining per nuclei ratio. changes in endothelial cell levels in the heart were determined from bulk RNAseq analysis using murine Microenvironment Cell Population counter (mMCP-counter) In all cases, data represent mean ± SD. Statistical significance was determined using Student’s unpaired t-test. \**p*<0.05.

Transcriptomic analysis of the hearts of R848-treated mice confirmed that resiquimod treatment was driving type I IFN responses, ischemia, endothelial remodeling, cardiomyocyte stress and active fibrosis in mice (Figure 2A, Supplemental Figure 3A), with a notable increase in *Nppa* and *Npr1* (markers of cardiac stress), and *Myl4* and *Myl7* (both fetal atrial genes reactivated during cardiac adaptation). Notably, a strong concordance was observed between RNA-Protein levels in the pathways regulating IFN, inflammation and IL6 signaling, all of which increased (Figure 2B), whereas genes and proteins relating to oxidative phosphorylation were decreased in R848-treated hearts (Figure 2C, Supplemental Figure 2A&B). We also noted a decrease in metabolites associated with the TCA cycle (Supplemental Figure 3C and Figure 2D, orange circles). Integration of the proteomic and metabolomic data using MixOmics, identified the strongest integrated enrichment (purple circles) for metabolic pathways (glucose, purine/pyrimidine, amino acid and fatty acid metabolism) (Figure 2D), supporting our evidence that metabolic changes are occurring in the R848-treated heart tissue.

**Figure 2:**
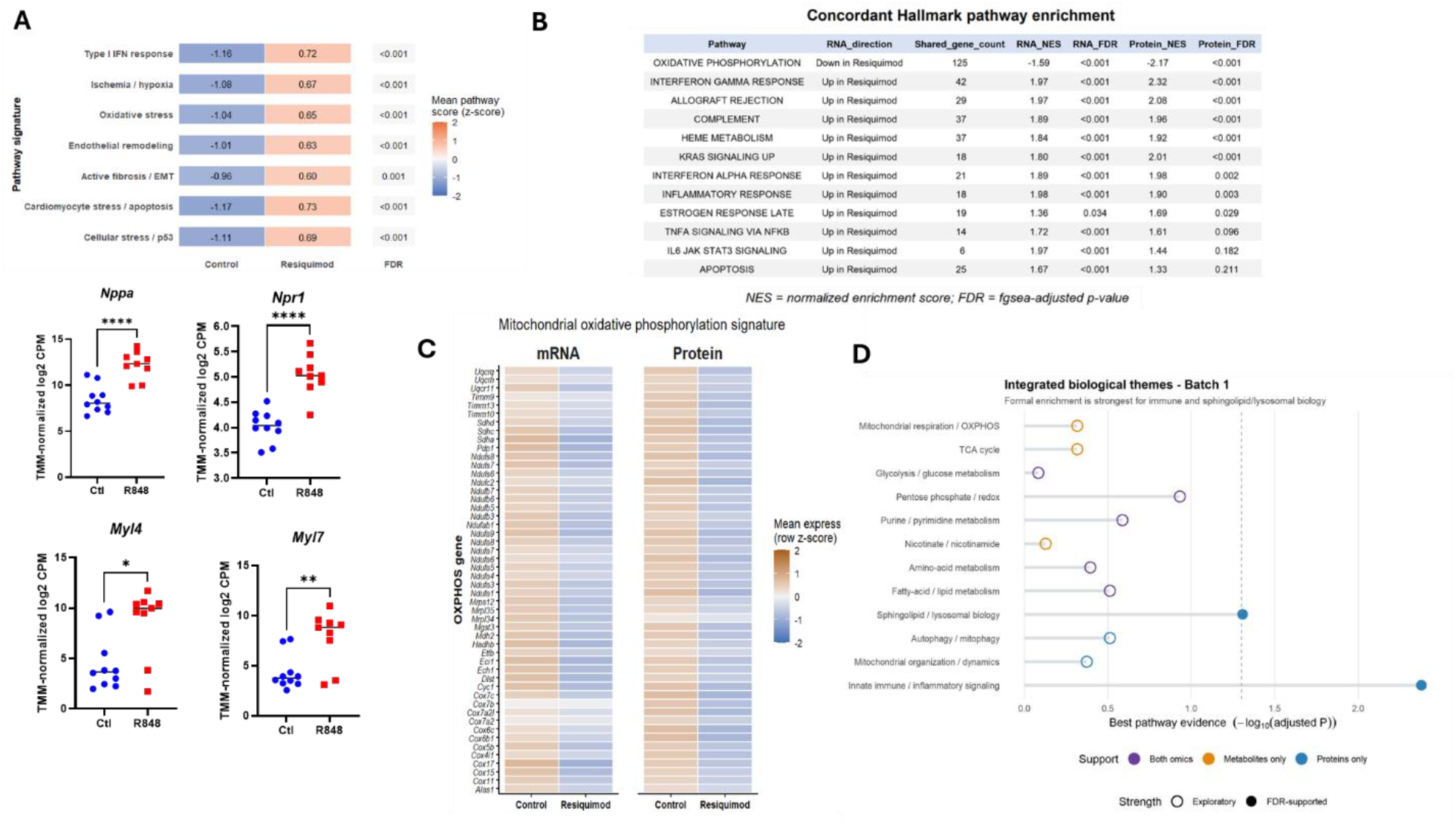
Resiquimod (R848) treatment induces genomic and proteomic changes, consistent with increased metabolic stress and cardiomyocyte remodeling. (A) Gene set variation analysis (GSVA) was carried out on bulk RNAseq from from hearts of control (n=5) and R848-treated mice (n=8). GVSA scores were calculated for MSigDB Hallmark gene sets using log_2_-transformed DESeq2-normalized RNA-seq counts. Scores were z-standardized across samples, and the heatmap shows mean pathway scores for hearts from control and R848-treated mice. Numbers within the tiles represent mean z-scores; FDR values were calculated from two-sided comparisons between groups using the Benjamini–Hochberg correction; (B) Ranked RNA-seq and protein fold-change statistics were independently analyzed using fgsea with the MSigDB Hallmark gene sets. Pathways with FDR < 0.25 in both datasets and concordant normalized enrichment score (NES) direction were considered shared. NES indicates the direction and magnitude of pathway enrichment, while FDR represents the fgsea-adjusted significance. Concordant genes were defined as unique genes within each Hallmark pathway showing the same direction of change in both RNA and protein datasets; (C) 50 OXPHOS genes with the strongest decreases in both RNA and protein were selected based on their combined RNA–protein fold changes. Heatmap values show mean row-scaled expression for Control and R848 heart samples; (D) Integrated proteomic and metabolomic pathway analysis of R848-treated mouse hearts. Biological pathways were assessed using matched proteomic and metabolomic datasets from control and R848-treated mice. Pathways are grouped by biological function, with colors indicating support from proteins only, metabolites only, or both datasets. Pathway evidence is represented as −log10(adjusted P value), with filled circles indicating FDR-supported enrichment (FDR < 0.05) and open circles indicating exploratory findings. The dashed line represents the FDR significance threshold.

Given that cardiomyocytes are the most energetically demanding cell type in the heart, and the cells most susceptible to stress and inflammation (18), we hypothesized that R848 treatment may be driving altered mitochondrial function in cardiomyocytes. We therefore conducted an *in vitro* analysis of the effects of acute and chronic IFN, using AC16 human ventricular cardiomyocytes (19). These cells exhibit key cardiac markers—contractile proteins, gap junction proteins, and transcription factors-indicative of cardiomyocyte identity and have been used to study mitochondrial metabolism and hypoxic signaling in response to hypoxia and reoxygenation (20). We structured our *in-vitro* model in a way that allowed us to study the effects of both acute and chronic exposure to type I IFN (Figure 3A). AC16 cells were stimulated with 1000ug/ml of IFNα for 24-hours. The cells were then washed twice with PBS to remove the IFNα and allowed to either rest for 4 days (primed) or rested (3 days) and then restimulated for another 24-hours post rest (chronic IFN). In addition, we treated cells with IFNα for 24-hours to assess the effects of acute IFNα treatment on the cells (acute IFN). All cells were seeded at the same time and harvested on the same day (Figure 3A). Using this treatment protocol we first looked at the effects of primed, acute or chronic IFN on gene expression in the AC16 cells. qPCR of RNA isolated from the different groups compared gene expression of ISGs such as MX1, IRF7, ISG15, IFI44, and IFI27 across the 4 groups (unstimulated, acute IFN, primed IFN and chronic IFN). ISGs were upregulated immediately after stimulation with IFNα (acute IFN), and in the primed + IFN group (chronic IFN) but this increase was not observed in the primed cells nor did priming increase the IFN response in the primed + IFN (chronic) group (Supplemental Figure 4). RNAseq analysis found that priming cells alone drove the expression of a small subset of ISGs (data not shown) whereas the greatest changes in gene expression were observed in the acute and chronically activated AC16s when compared to control. As shown in the Venn diagram (Supplemental Figure 5A), the majority of genes were common between both and gene set variation analysis showed that the key pathways targeted were IFNα and IFNγ responses (Supplemental Figure 5B). However, GSVA showed that priming with IFNα drove changes in genes associated with oxidative phosphorylation, glycolysis and ROS production, indicating that although the majority of changes were in response to acute IFN stimulation, chronic IFN exposure may drive mitochondrial remodeling (Supplemental Figure 5C).

To measure if there was a change in metabolic activity of these cells due to acute and chronic IFN exposure, we conducted mitochondrial and glycolytic stress assays using Seahorse Extracellular Flux Analysis analyzer and observed that primed and chronic exposure groups show impaired oxidative phosphorylation and glycolysis, and decreased basal respiration, maximal respiration, ATP production (Figure 3B-C), indicating mitochondrial dysfunction. In line with this, priming or chronically exposing cells to IFN triggered loss of mitochondrial membrane potential (Figure 3D), potentially contributing to the changes in metabolism seen. In addition, we observed a significant increase in mitochondrial proteins in the primed and chronic groups compared to either control or acute IFNα stimulated cells, using target mass spectrometry analysis (Figure 3E). These changes in mitochondrial proteins spanned different mitochondrial pathways such as TCA cycle (SDHA, MDHM, ACON), electron transport chain (SODM), mitochondrial membrane transport (TIM50, COX2), mitochondrial transcription (TFAM) and fission (DRP1) (Supplemental Figure 4). Proteomic analysis of total cellular proteins confirmed that priming drove increases in proteins associated with mitochondrial and metabolic pathways. Integrating the proteomic and transcriptomic datasets confirmed a coordinated increase in genes and proteins associated with mitochondrial respiration, oxidative phosphorylation and mitochondrial translation in cells that had been primed with IFN for 4 days (Figure 3F). Similarly, priming also increased genes and proteins regulating mitochondrial organization and dynamics. Combined our data strongly suggests that the cells are attempting to adapt or compensate for stress due to chronic IFN exposure by increasing the abundance of mitochondrial machinery, to attempt to mitigate defects in mitochondrial function or integrity that could ultimately be limiting both aerobic and anaerobic ATP generation.

Mitochondrial depolarization, reduced oxidative phosphorylation (OxPhos) and extracellular acidification rate (ECAR), accompanied by an increase in mitochondrial protein expression, suggested to us that priming/chronic IFN exposure was triggering severe mitochondrial dysfunction and loss of membrane potential and potentially a compensatory biogenesis response. Using MitoTracker Green, we observed that mitochondrial mass increased when cells were primed or chronically exposed to IFN (Figure 4A), in addition we observed a greater degree of fragmentation of mitochondria in response to IFN treatment (Figure 4B & Supplemental Figure 6), characterized by a decrease in branches per mitochondria, junctions, junctions, triple point and branch length per mitochondria (Supplemental Figure 6). This was consistent with the observed increase we saw in Drp1 at the transcriptional level (Figure 4C). Expression of mitophagy components LC3II/LC3I and p62 also increased, consistent with increased efforts to remove defective mitochondria under conditions of stress (Figure 4D). Importantly, recombinant asporin (rASPN), a critical protector of mitochondrial integrity, was able to rescue mitochondrial branch length in AC16 cells chronically exposed to IFN (Figure 4E).

**Figure 3:**
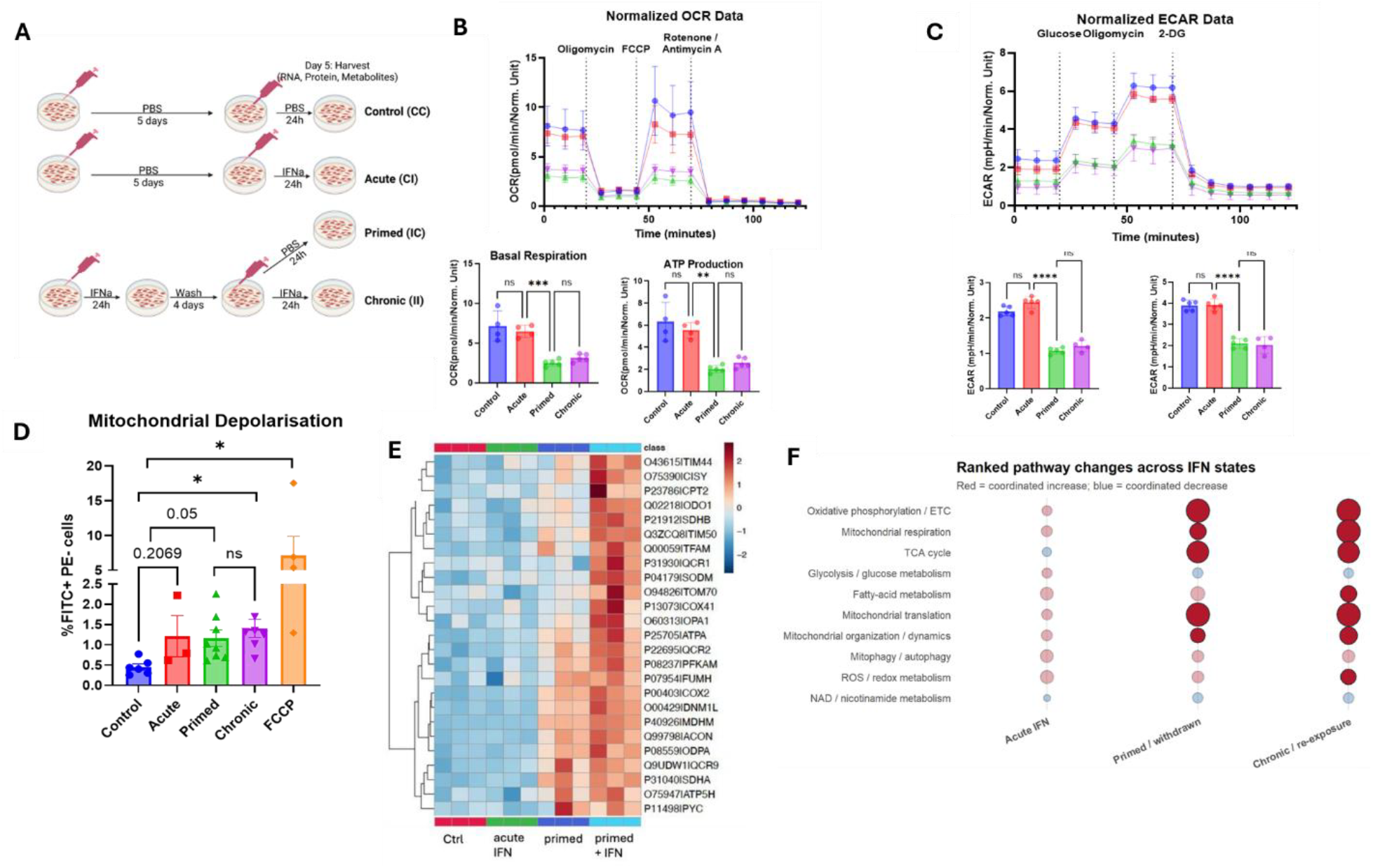
Chronic IFN exposure drives mitochondrial dysfunction and depolarization in AC16 cardiomyocytes. (A) Schematic showing AC16 treatment and generation of 4 groups (n=3 for each) Control (CC), primed (IC), Acute IFN (CI) and Chronic IFN (II); (B) Seahorse Extracellular Flux Analysis (EFA) was used to monitor (B) oxygen consumption rates and (C) extracellular acidification rate (ECAR) in real time. Data represent mean ± SD of reads n=5 samples from one experiment, and are representative of data (A-D) n=5; (D) Mitochondrial depolarization in response to IFN treatment (n=3 per group) was assessed across our samples by flow cytometry using the fluorescent dye JC-1; (E) Proteins across our 4 groups (n-=3 per group) were isolated for targeted tandem mass spectrometry to measure 37 proteins associated with mitochondrial metabolism and function. A heatmap of the differentially expressed proteins across the 4 groups was generated using MetaboAnalyst; (F) Gene set variation analysis (GSVA) was performed using log_2_-transformed DESeq2-normalized counts. Bubble plots show changes in selected mitochondrial and metabolic pathways relative to control, with color indicating the direction of change (red, increased; blue, decreased) and bubble size representing the magnitude of pathway alteration.

**Figure 4.**
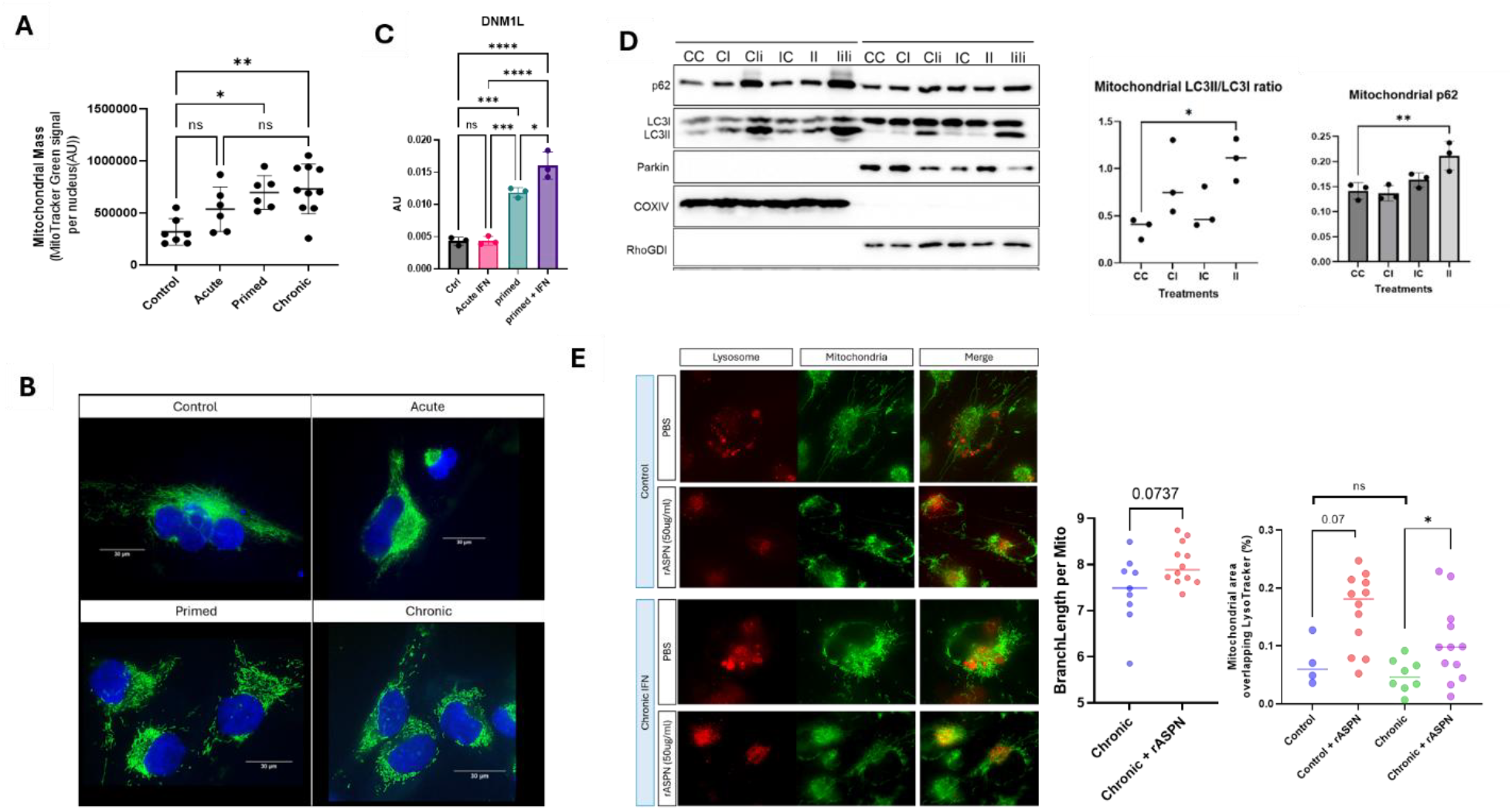
Chronic type I interferon results in increased mitochondrial mass, increased fragmentation and impairs mitochondrial quality control. (A) MitoTracker Green, a cell-permeant fluorescent dye that stains mitochondria in live cells was used to stain the four groups of AC16 cells. Mitochondrial mass was estimated by quantifying MitoTracker Green fluorescence intensity normalized to the number of nuclei using Fiji. Data are presented as mean ± SD, with individual points representing independent measurements. Statistical significance is indicated as ns, not significant; P < 0.05, P < 0.01.(B) Mitochondrial imaging shows progressive remodeling and fragmentation/simplification of the mitochondrial network across IFN states (C)Targeted measurement of DNM1 by MS/MS, A.U.=Arbitary Units; (D) Western blotting for p62, Parkin, COXIV, RhoGDI, LC3I and LC3III showed increased mitochondrial p62 levels and increased mitochondrial LC3III/LC3I ration (n=3 per group; CC=control, CI=Acute, IC= Primed, II+ Chronic). (e) AC16 human cardiomyocytes were cultured under control (PBS) or chronic interferon (IFN) conditions in the presence or absence of recombinant asporin (rASPN; 50 µg/mL). Representative fluorescence microscopy images show mitochondria (green), lysosomes (red), and merged channels. Mitochondrial–lysosomal colocalization was quantified using automated image analysis of independently segmented mitochondrial and LysoTracker-positive regions and expressed as the percentage of total mitochondrial area overlapping the LysoTracker-positive mask. Individual data points represent analyzed image fields.

When mitochondria are stressed or become depolarized, they release mitochondrial (mt) nucleic acids (mtRNA/DNA) into the cytosol. These can then activate RNA sensing (RIG-I, IFIH1) and DNA sensing (cGAS-STING) pathways to increase IFN and inflammatory gene expression or to enhance cell death and stress. In line with our data showing mitochondrial depolarization in chronically activated or primed AC16 cells, we observed that priming or chronically exposing AC16 cells to IFN showed increased release of mtDloop DNA into the cytosol (Figure 4A). Interestingly, this increase was accompanied by an increase in *STING1* expression under the same conditions (Figure 4B). Importantly, the cGAS inhibitor, G140, inhibited ISG (IFIT1 and ISG15) expression in primed and chronically IFN exposed cells, whereas not in the acute IFNa treated samples (Supplemental Figure 7). RNAseq analysis of chronically treated AC16s pretreated wit or without G140 confirmed these results (Figure 4C) and also identified key pathway responses that were controlled by G140, namely hypoxia, glycolysis and OXPHOS, whereas pathways related to cell cycle regulation were turned on in G140 treated cells, indicating a rescue of phenotype (Figure 5D). Translating these findings back to our resiquimod model, we observe increases in cGAS, TBK1 and IRF3 at the gene and protein level, in addition to increased IFN stimulated genes/proteins, increase in autophagy and mitophagy machinery and a decrease in genes and proteins associated with mitochondrial function (Figure 5E). These findings have important implications for autoimmune driven CVD, specifically implicating the IFN system and cytosolic DNA sensing pathways in altering mitochondrial dynamics and ultimately cardiomyocyte function.

**Figure 5:**
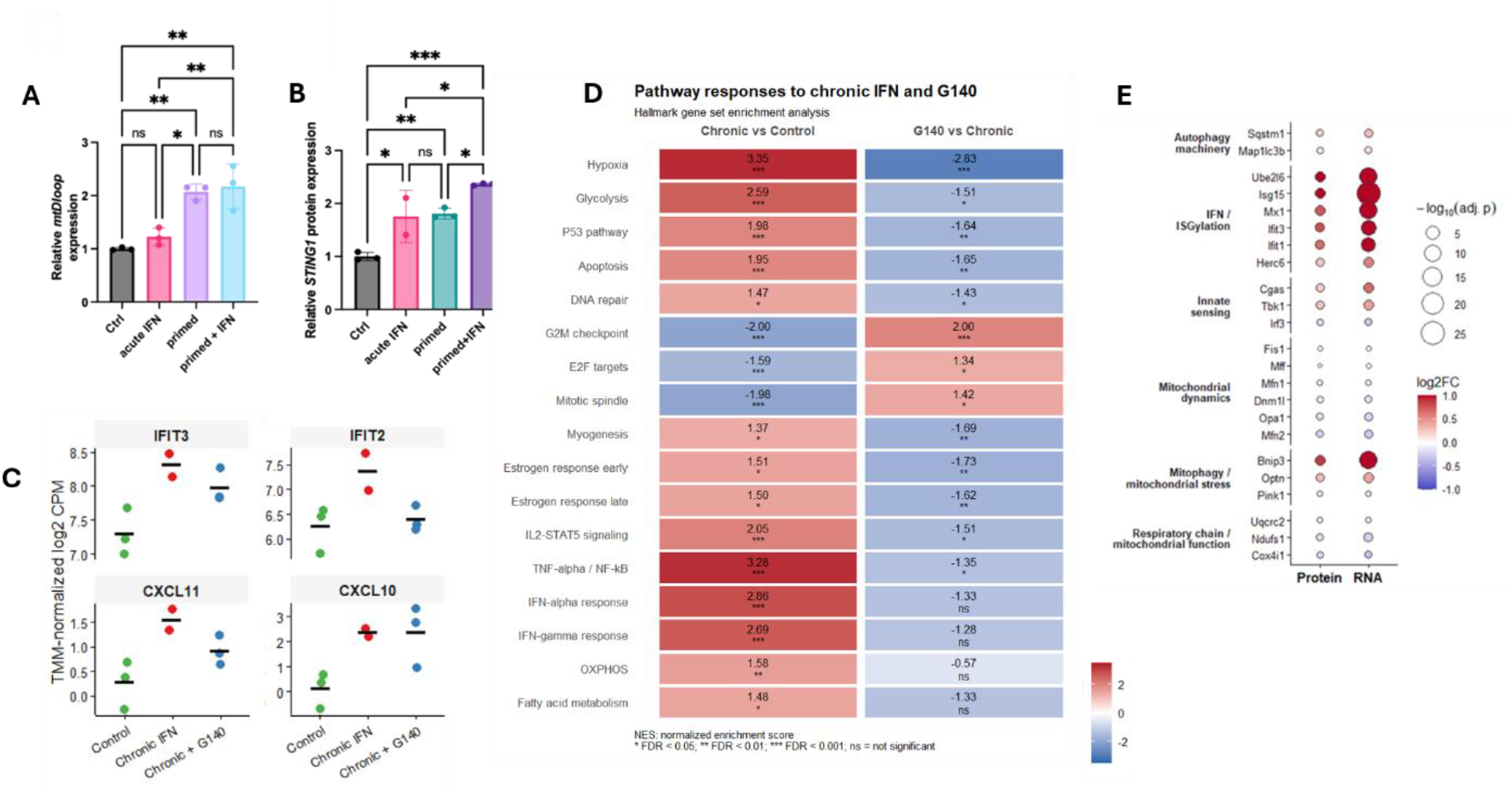
Mitochondrial damage promotes mtDNA release and activates a cGAS–STING-dependent inflammatory feed-forward loop. (A) Cytosolic mtDNA was isolated from the four groups of AC16 cells and PCR for expression *mtDloop* used to determine release. (B) mRNA was isolated from whole cell lysates from treated AC16 cells and qPCR was used to determine *STING1* expression. Statistical significance was determined using One-Way ANOVA and Bonferroni’s multiple comparison test. \**p*<0.05, \*\**p*<0.01, \*\*\**p*<0.001 (C-D) Bulk RNAseq was carried out on RNA from control (n=3) and chronic IFN-treated with or without G140 AC16 cells (n=3 per group). (C) shows gene expression changes on representative cGAS-driven genes; (D) Gene set variation analysis (GSVA) was carried out on bulk RNAseq from control (n=3) and chronic IFN-treated with or without G140 AC16 cells (n=3 per group). GVSA scores were calculated for MSigDB Hallmark gene sets using log_2_-transformed DESeq2-normalized RNA-seq counts. Scores were z-standardized across samples, and the heatmap shows mean pathway scores for Chronic vs control and G140 vs Chronic cells (n=3 per group). Numbers within the tiles represent mean z-scores; FDR values were calculated from two-sided comparisons between groups using the Benjamini–Hochberg correction. In all cases, data represent mean ± SD of reads from triplicate samples. (E) Bubble plot showing changes in selected proteins and corresponding RNA expression between control and resiquimod-treated mouse hearts. Genes are grouped by biological function. Bubble color represents log2 fold change (RES vs. control), while bubble size indicates statistical significance (−log10 adjusted P value).

## Discussion

Hasham *et al* showed that epicutaneous application of the TLR7/8 agonist R848, resiquimod, to CFN hybrid mice (cross of C57Bl6/J, FVB/NJ, and NOD/ShiLtJ) led to development of SLE-like autoimmune disease, characterized by the overproduction of type I interferons and IFN-stimulated genes. Importantly, the mice showed rapid progression from acute myocarditis to severe dilated cardiomyopathy and histopathology revealed widespread heart inflammation resembling autoimmune pancarditis (17).

Systemic effects of type I interferons on heart function was assessed using the resiquimod mouse model and both ejection fraction and fractional shortening were reduced which indicates a reduction in cardiac output due to possible left ventricular dysfunction. We also observed cellular hypertrophy in the heart sections from the resiquimod-treated mice using histology. This suggests an adaptive but pathological response of cardiomyocytes to ongoing stress, linking the immune activation to adverse structural remodeling. Transcriptomic and proteomic profiling of the heart tissue establishes the resiquimod treatment resulted in an enrichment of interferon-driven immune pathways such as T cell responses and myeloid differentiation. Simultaneously, we observed a suppression of fundamental mitochondrial metabolic processes including the TCA cycle and electron transport chain and the mechanistic connection between the two could indicate a shift from oxidative energy metabolism toward immunometabolic reprogramming in the resiquimod-treated mouse hearts. Metabolomic analysis further reveals accumulation of ischemia-associated intermediates such as succinic acid, deoxyinosine and pyridoxic acid, and pathway enrichment analysis highlights dysregulation of purine metabolism, pentose phosphate pathway, and the malate-aspartate shuttle. These results suggest that systemic type I interferon exposure can remodel cardiac function and metabolism and implicates cross-talk between immune activation, mitochondrial energy failure, and metabolite signaling as key contributors to interferon-driven heart disease. In response to ischemic injury, metabolic stress, or systemic inflammation, immune cells such as macrophages, T cells, B cells, and neutrophils infiltrate the myocardium and undergo shifts in core metabolic pathways by moving between oxidative phosphorylation and glycolysis according to their activation and functional state. Metabolic imbalances themselves can also induce systemic inflammation, which disrupts heart structure and function by promoting maladaptive remodeling through cytokines, excess metabolic substrates, and paracrine signaling (21). These shared inflammatory pathways in cardiac cells cause progressive damage, leading to metabolic cardiomyopathy and, clinically, to heart failure with preserved ejection fraction (HFpEF)(21). Obesity and diabetes share cellular changes resembling those seen in HFpEF, such as oxidative stress, mitochondrial dysfunction, lipotoxicity, hypertrophy, ECM buildup, and microvascular disease (22).

One highly novel finding from our metabolomics data was the observation of a dramatic increase in taurocholic acid levels, a bile acid, in the hearts of resiquimod treated mice. Bile acid metabolism is strongly linked to cardiovascular diseases (CVDs) and metabolic disorders either by directly interacting with heart muscle cells and modifying contraction and electrical conduction or indirectly by modulating metabolic processes such as heart function regulation, cholesterol metabolism, and atherosclerotic plaque development (23, 24). TCA specifically can induce arythmias and contractile dysfunction by altering calcium ion dynamics in cardiomyocytes (25). Other studies have linked excess bile acids with reduced fatty acid metabolism in cardiomyocytes, directly linking bile acids to reduced cardiomyocyte function (24). In SLE, patients have been shown to have elevated TCA in their feces, potentially driven by an imbalance in the gut microbiome (26). However, to date no link between TCA and SLE heart involvement has been suggested, and ours is the first study to identify an IFN-driven increase in heart tissue in resiquimod treated mice. Although more work needs to be done definitively establish it, our findings suggest a novel link between bile acid metabolism and cardiac dysfunction in the context of chronic interferon-driven disease.

Our multiomic analysis of the heart tissue from resiquimod treated mice strongly implicated cardiomyocytes as the cell type undergoing pathologic changes in the heart. Transcriptomic analyses showed that both acute and chronic IFNα elevated ISG expression similarly, which is in line with literature reporting strong ISG induction in cardiomyocytes following type I interferon signaling and myocardial injury (7). Notably, chronic IFNα exposure led to distinct upregulation of genes related to cell migration and metabolic regulation, while primed and control samples showed top enrichment in ISGs and downregulation of genes involved in cholesterol biosynthesis. Recent studies have shown Toll-like receptors and interferons as mediators of the reprogramming of cholesterol homeostasis and regulation of sterol metabolic pathways in immune cells has been as acknowledged an integral response to host defense (27, 28).

At the proteomic level, primed cells compared to controls exhibited enrichment of pathways such as ‘mitochondrial protein degradation’, ‘mitochondrial translation’, and ‘respiration and respiratory electron transport’. Consistent upregulation of ISGs was seen with both acute and chronic IFNα exposure, but GSEA revealed features unique to the chronic IFN exposure that overlapped with the pathways observed in the primed cells, hence IFN priming contributed to those differentiating pathways. One study in Chaga’s disease observed that AC16 cells co-exposed to IFN-γ and TNF showed protein and gene expression changes associated with mitochondrial dysfunction, altered fatty acid metabolism and cardiac cell death (29). IFNα not only induces the expression of RIG-I and cGAS, but these sensors also detect cytosolic nucleic acids such as mitochondrial DNA released during stress or injury and trigger further type I interferon production (30-32). Both the cGAS-STING and RIG-I pathways participate in cardiac injury and remodeling by promoting chronic inflammation and innate immune activation in the heart, forming a feed-forward loop that can exacerbate myocardial damage (33, 34). Our findings are novel in showing that IFNα priming changes the functional responses of human ventricular cardiomyocytes beyond ISG induction, particularly impacting mitochondrial organization, protein turnover, and cholesterol metabolism. This may provide new mechanistic insight into how chronic IFN exposure shapes cardiomyocyte function and contributes to the heart defects observed in lupus and interferon-driven heart disease models.

In line with mitochondrial effects of chronic IFNa, Mitostress and Glycolysis stress assays showed that chronic IFN exposure resulted in decrease in both oxidative phosphorylation and glycolysis. This is interesting since during heart disease and cellular stress, cardiomyocytes undergo metabolic reprogramming, shifting from reliance on oxidative phosphorylation (OXPHOS) toward increased glycolysis to meet energy demands (35). This switch promotes cell survival under hypoxic or ischemic conditions but contributes to impaired contractile function and disease progression in the failing heart (36). The dual metabolic suppression of OXPHOS and glycolysis reflects a loss of metabolic flexibility and an inability to compensate for energy deficits through glycolysis (37). This is observed in advanced heart failure, drug-induced cardiotoxicity, or extreme metabolic stress, where mitochondrial function is severely impaired and cellular energy generation is stalled (38, 39). A targeted mass spectrometry assay for 34 mitochondrial proteins shows that chronic IFN also increased many mitochondrial proteins involved in the TCA cycle (SDHA, MDHM, ACON), electron transport chain (SODM), mitochondrial membrane transport (TIM50, COX2), mitochondrial transcription (TFAM) and fission (DRP1) (40). This suggests that chronic IFN signaling drives compensatory adaptations in cardiomyocytes, enhancing mitochondrial content and machinery to meet increased energy and metabolic demands or counteract stress.

The cGAS-STING and RIG-I signaling pathways are critical mediators of innate immune responses triggered by mitochondrial DNA (mtDNA). Cardiac stress, injury, or mitochondrial dysfunction can lead to the escape of mtDNA into the cytosol, where it is sensed by cGAS, activating the STING pathway and promoting a cascade of inflammatory responses, including increased production of type I interferons and pro-inflammatory cytokines (33). This heightened inflammation has been shown to drive pathological cardiac remodeling, hypertrophy, fibrosis, and dysfunction, and pharmacologic or genetic inhibition of cGAS-STING signaling can ameliorate these effects in experimental models (41). Similarly, RIG-I, another key cytosolic sensor, detects nucleic acids (including mtRNA) and orchestrates inflammatory responses in cardiac fibroblasts by increasing the production of cytokines such as IL-6 and IL-8, contributing to endothelial and myocardial injury, and ultimately advancing cardiac pathology (42). Together, these innate sensing pathways represent crucial molecular links between mitochondrial damage, persistent inflammation, and the onset and progression of heart disease. Our proteomics analysis revealed that both acute and chronic IFNα exposure lead to elevated levels of STING1 and RIG-I, with STING1 remaining persistently high in the primed group compared to control cells. To further investigate, we assessed cytosolic mtDNA release by isolating the cytosolic fraction from AC16 cardiomyocytes and we found that the primed and primed + IFN groups exhibited a marked increase in these mitochondrial DNA sequences in the cytosol, demonstrating that prolonged IFNα exposure promotes substantial mtDNA release in cardiomyocytes. While this observation is in line with literature showing that cGAS-STING and RIG-I pathways can be activated by mtDNA release during cardiac stress, driving type I interferon signaling and inflammation, persistent STING1 elevation and robust mtDNA release in cardiomyocytes represents a novel mechanistic insight into how systemic interferon signaling may drive heart disease in SLE and present targets for therapeutic intervention.

