## Supplemental FIgures for "Mitochondrial instability contributes to IFN-driven heart disease"

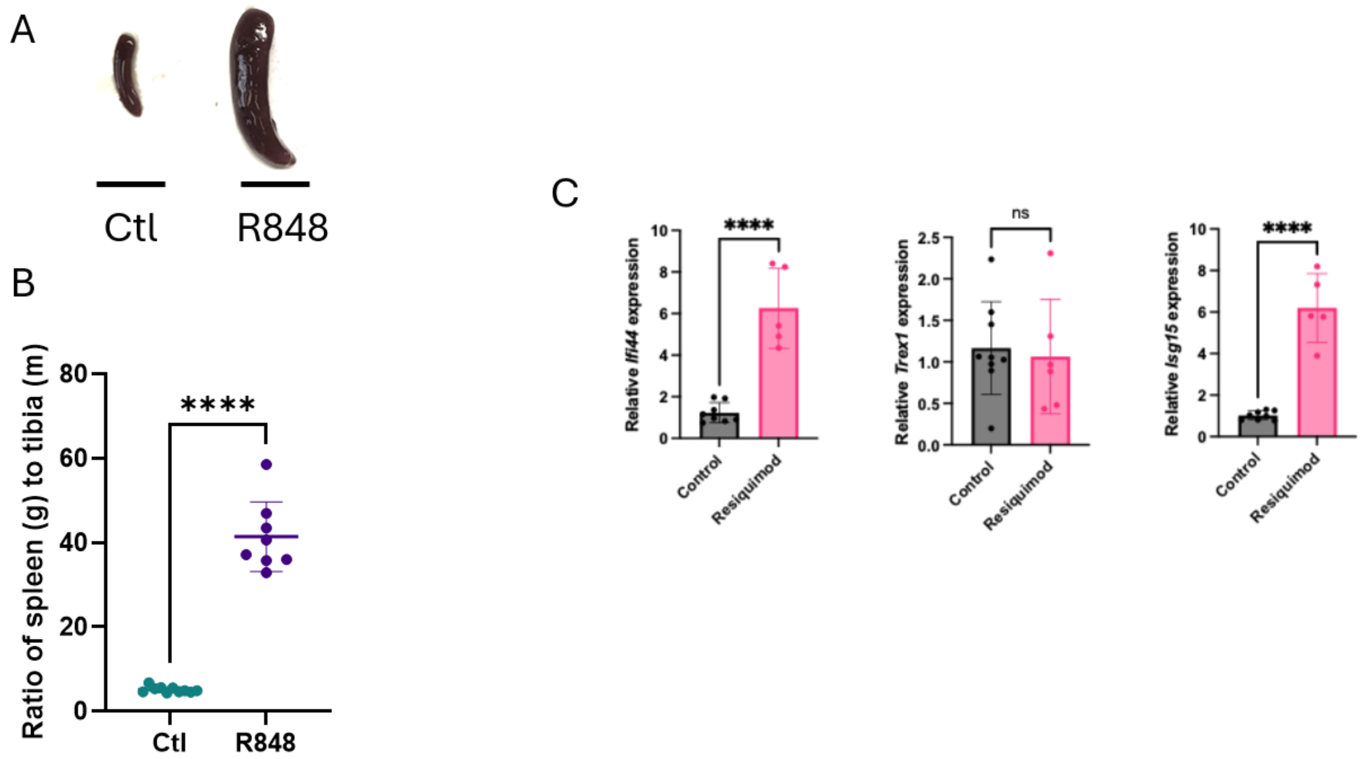


**Supplemental Figure 1 Resiquimod treated mice showed systemic inflammation through splenomegaly and increase in ISG gene expression in splenocytes.** (A) Representative image of spleens of control mice (left) and R848-treated mice (right) (B) Spleen weight normalized to tibia length; (C) Gene expression measured through qPCR of RNA extracted from splenocytes. Control and treated mice grouped as there were no changes between sex differences. Data represent mean ± SD. Statistical significance in B was determined using One-Way ANOVA and Bonferonni's 's multiple comparison test. *$p$<0.05, **$p$<0.01, ***$p$<0.001, ****$p$<0.0001. Statistical significance in C was determined using Student's unpaired t-test. *$p$<0.05, **$p$<0.01, ***$p$<0.001, ****$p$<0.0001.

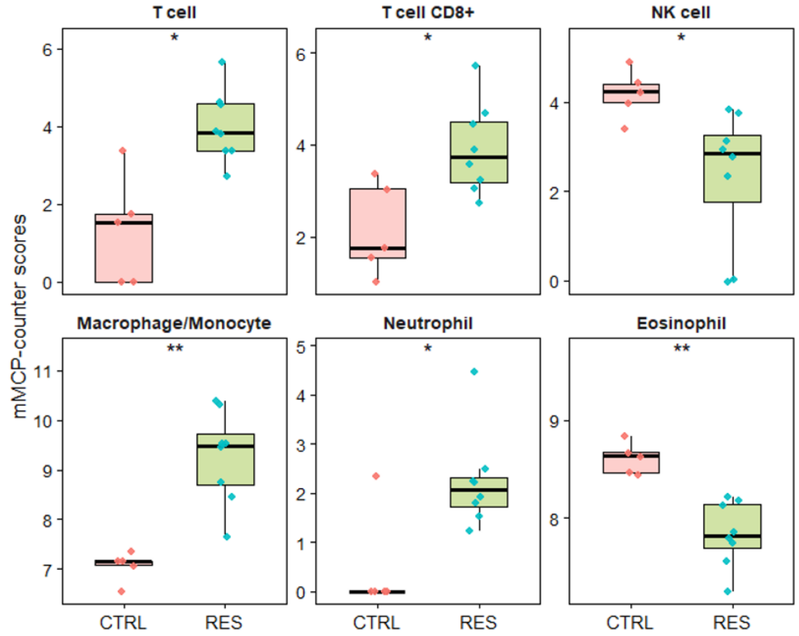


**Supplemental Figure 2. Resiquimod treatment alters immune cell populations in mouse hearts.** Immune cell populations were estimated from bulk RNA-seq data using mMCP-counter. Boxplots show relative abundance scores for the indicated cell populations in control (CTRL) and resiquimod-treated (RES) hearts. Each point represents an individual mouse. Statistical significance was assessed using Wilcoxon rank-sum tests (*P* < 0.05, **P** < 0.01).

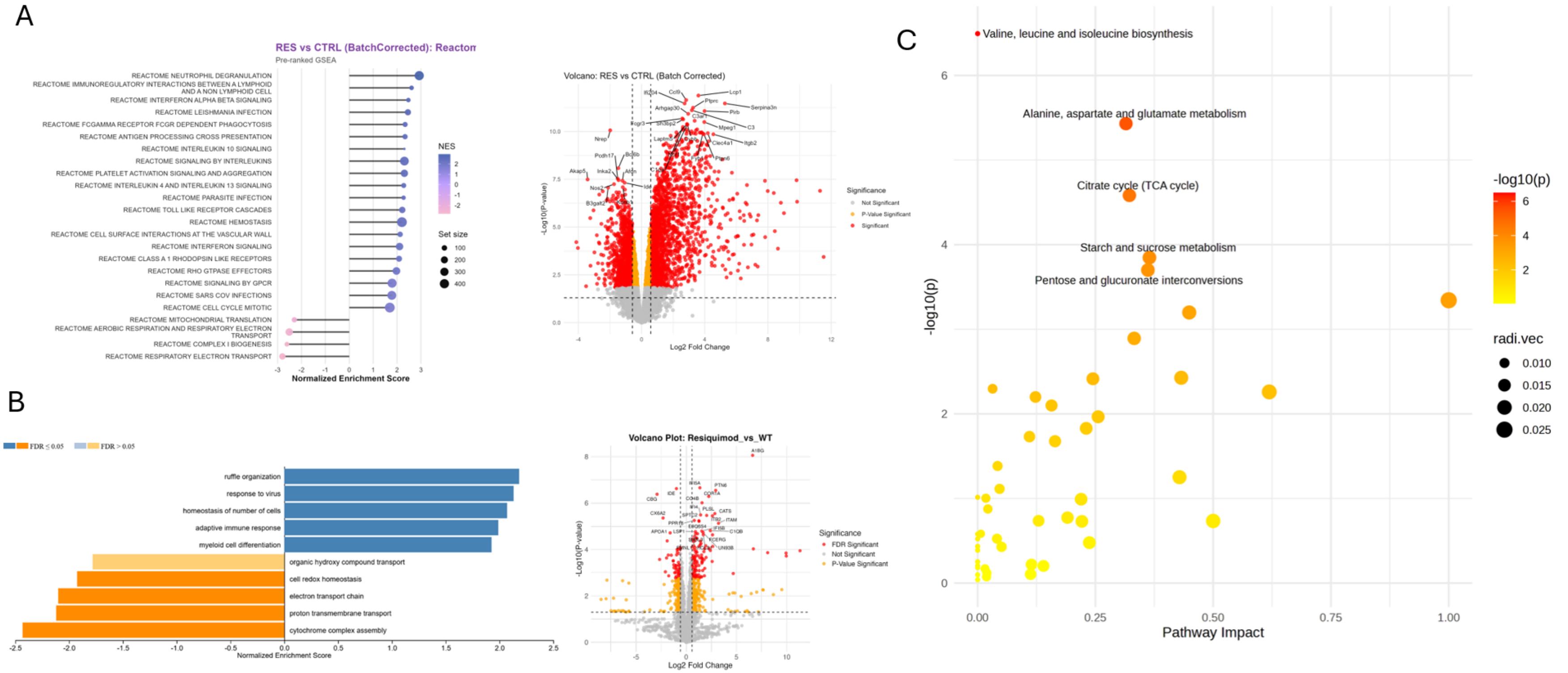


**Supplemental Figure 3. Resiquimod-treatment results in transcriptomic and proteomic changes consistent with altered immune response and altered TCA cycle in the heart**

(A) RNA and (B) proteins were extracted from hearts of control and R848-treated mice and analyzed by RNAseq and Mass spectrometry, respectively (control n=10, R848-treated n= 9) (A, B) Gene set enrichment analysis (GSEA) was performed against the Reactome pathway database (version v2025.1) from MSigDB. Significance was assessed using 1,000 permutations, and the top-ranking pathways were identified based on the normalized enrichment score (NES) and a false discovery rate (FDR) q-value < 0.25 and are represented here. Right hand panel of (A) and (B) showing volcano with top genes highlighted; (C) Scatter diagram summarizing pathway enrichment by pathway impact (x-axis) and enrichment significance (y-axis, -log10 p-value) of differentially expressed metabolites (>2 fold change, false discovery rate (FDR) q-value < 0.25).

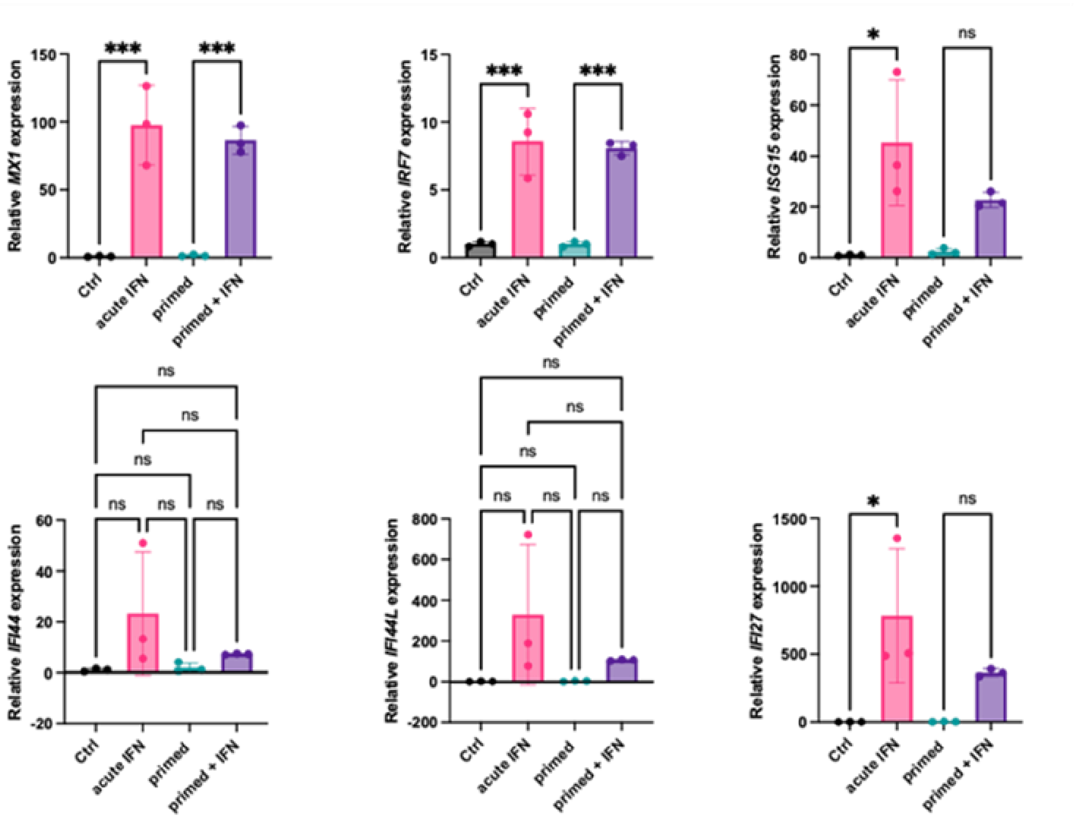


**Supplemental Figure 4. IFNα increases expression of ISGs in AC16 cardiomyocytes**

AC16 cells stimulated as described in Figure 5A. RNA was extracted from the samples and qPCR was conducted to measure ISG expression values. Each group had triplicates, and statistical analysis was determined by One-way ANOVA and Bonferroni's post hoc analysis for multiple comparisons. $^{**}p<0.01$, $^{***}p<0.001$, $^{****}p<0.0001$.

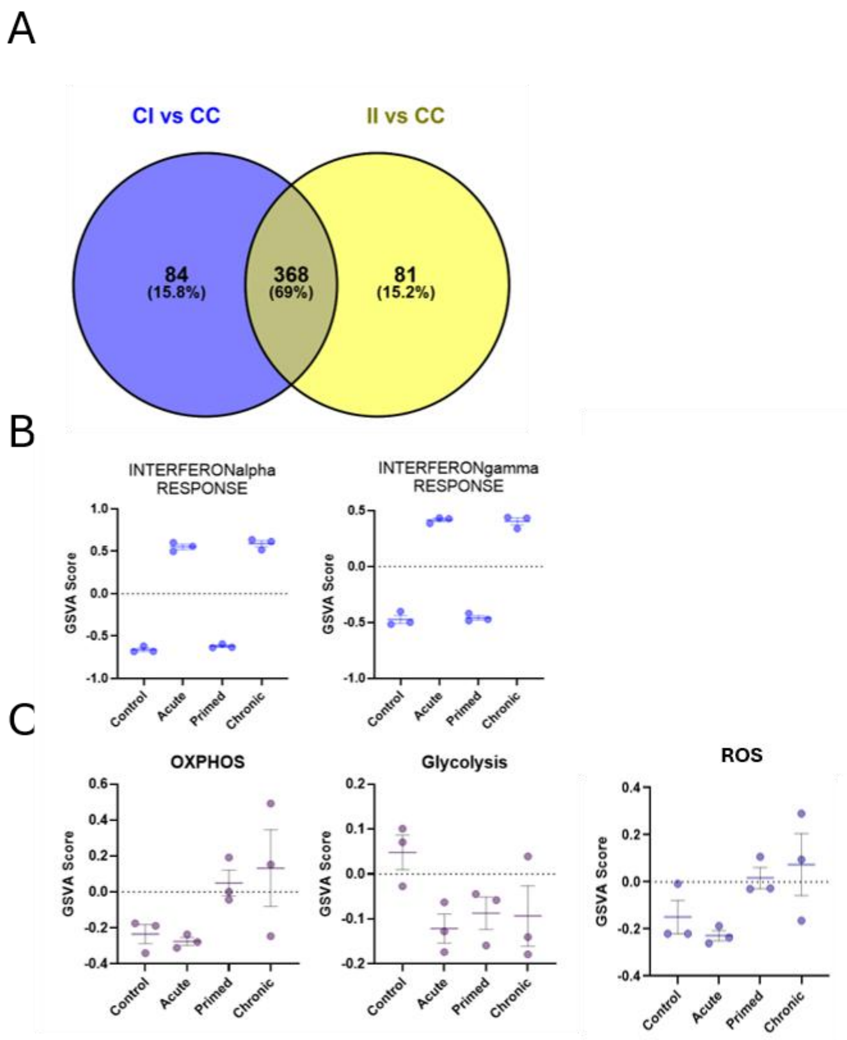


**Supplemental Figure 5. Acute and chronic IFNα exposure in AC16 cardiomyocytes resulted in differential changes in gene expression**

(A) Venn diagram showing the numbers of common and unique DEGs between CI vs CC and II vs CC; (B)

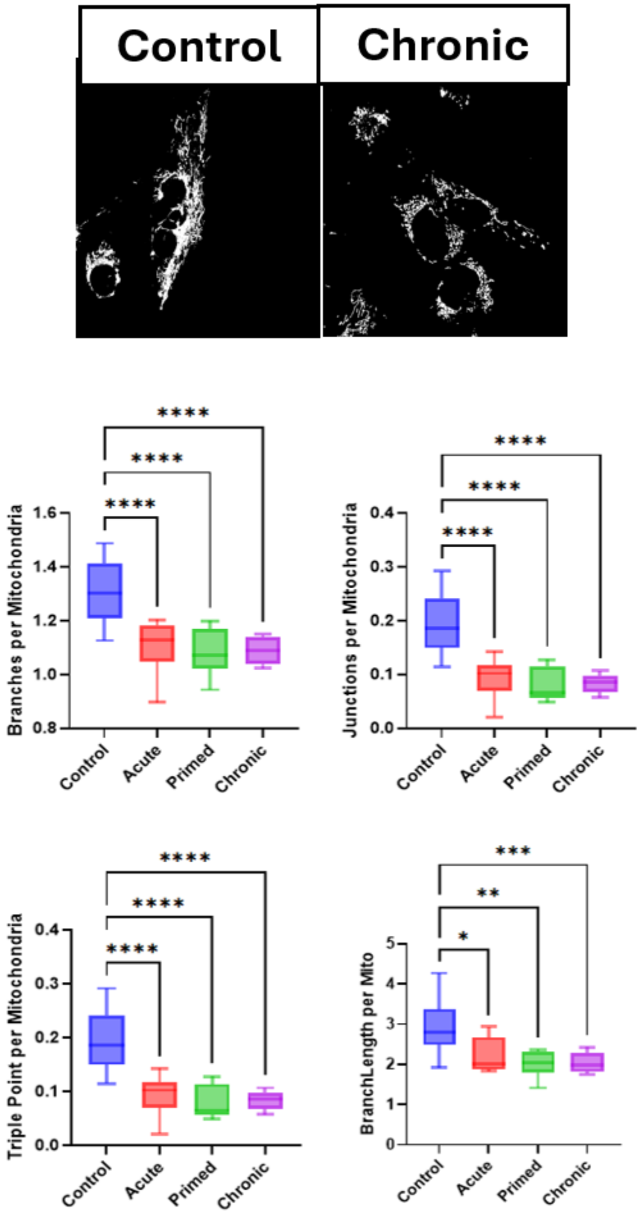


**Supplemental Figure 6. Altered mitochondrial morphology following IFN treatment in AC16 cardiomyocytes.**
Representative mitochondrial masks used for morphological analysis in Fiji (top). Mitochondrial morphology was quantified across control, acute, primed, and chronic IFN treatment conditions, assessing branches, junctions, triple points, and branch length per mitochondrion. Boxplots show median and interquartile range. Statistical significance is indicated as *p<0.05, ***p*<0.01, ****p*<0.001, ****p<0.0001.

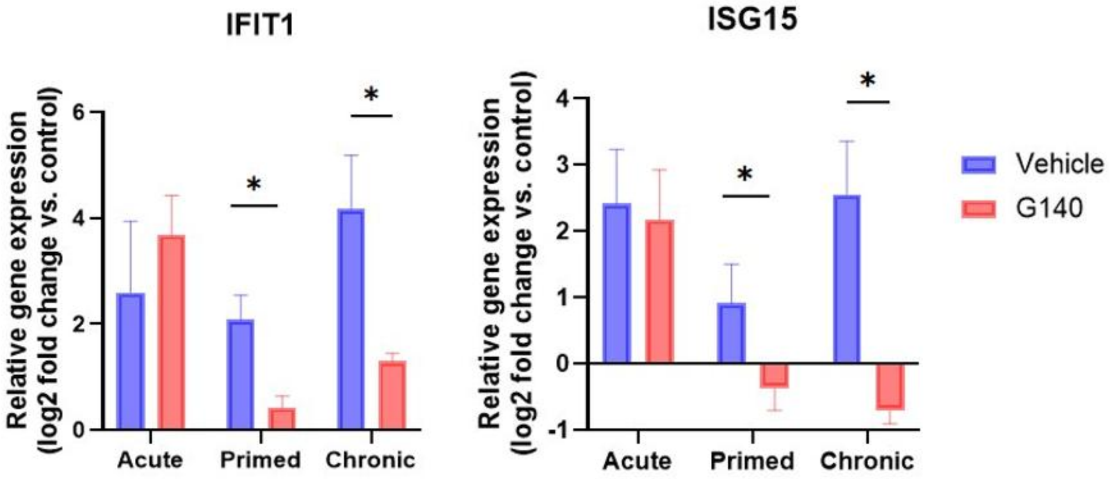


**Supplemental Figure 7: G140 inhibits chronic IFN effects but not acute in AC16 cells.** AC16 cells were treated with IFN as indicated with or without 1 hour pretreatment with cGAS inhibitor G140. mRNA was isolated from whole cell lysates from treated AC16 cells and qPCR was used to determine *IFIT1* and *ISG15* expression. In all cases, data represent mean ± SD of reads from triplicate samples. In all cases, statistical significance was determined using One-Way ANOVA and Bonferroni's multiple comparison test. *$p$<0.05, **$p$<0.01, ***$p$<0.001
